# Developing and deploying an integrated workshop curriculum teaching computational skills for reproducible research

**DOI:** 10.1101/2021.06.15.448091

**Authors:** Zena Lapp, Kelly L. Sovacool, Nick Lesniak, Dana King, Catherine Barnier, Matthew Flickinger, Jule Krüger, Courtney R. Armour, Maya M. Lapp, Jason Tallant, Rucheng Diao, Morgan Oneka, Sarah Tomkovich, Jacqueline Moltzau Anderson, Sarah K. Lucas, Patrick D. Schloss

**Author notes:** These authors contributed equally to this work (Zena Lapp, Kelly L. Sovacool). Correspondence: Patrick D. Schloss < >.

## Abstract

Inspired by well-established material and pedagogy provided by The Carpentries (Wilson 2016), we developed a two-day workshop curriculum that teaches introductory R programming for managing, analyzing, plotting and reporting data using packages from the tidyverse (Wickham et al. 2019), the Unix shell, version control with git, and GitHub. While the official Software Carpentry curriculum is comprehensive, we found that it contains too much content for a two-day workshop. We also felt that the independent nature of the lessons left learners confused about how to integrate the newly acquired programming skills in their own work. Thus, we developed a new curriculum that aims to teach novices how to implement reproducible research principles in their own data analysis. The curriculum integrates live coding lessons with individual-level and group-based practice exercises, and also serves as a succinct resource that learners can reference both during and after the workshop. Moreover, it lowers the entry barrier for new instructors as they do not have to develop their own teaching materials or sift through extensive content. We developed this curriculum during a two-day sprint, successfully used it to host a two-day virtual workshop with almost 40 participants, and updated the material based on instructor and learner feedback. We hope that our new curriculum will prove useful to future instructors interested in teaching workshops with similar learning objectives.

## Statement of Need

For the past five years, the University of Michigan instance of The Carpentries has taught workshops using versions of curriculum originally created by The Carpentries organization. In that time, our instructors found several advantages and disadvantages to using the original Software Carpentry curriculum. Some of the advantages were that any programming language lesson (e.g., R or Python) could be paired with lessons on the Unix shell and version control, lessons had been refined by many contributors over the years and taught at workshops around the world, and the instructional design demonstrated good pedagogy for teaching novice data science practitioners. However, The Carpentries materials have evolved from lesson plans to reference materials, and thus there was too much content for the time available during a two-day workshop. As a result, workshops taught with this material were inconsistent depending on who was teaching, and new instructors faced an overwhelming amount of work to prepare for their first workshop. Furthermore, the modular nature of the curriculum meant that each lesson was independent from the others, so it was not apparent to learners how all of the skills could be integrated for the purpose of a reproducible research project.

Given these constraints, we sought to create a new curriculum that would allow us to teach computational skills in an integrated manner, demonstrate the reproducible research workflows we use in our own work, deliver an appropriate and consistent amount of content, and reduce the burden for new instructors to get involved, all while maintaining the same inclusive pedagogy that has been refined by The Carpentries organization.

## Collaborative Curriculum Development

We drew on the expertise of The Carpentries community at the University of Michigan to develop a custom curriculum that would meet our goals (Figure 1). To start, we organized a two-day sprint, where members of our community worked collaboratively to create an initial draft of the content. During the sprint, we met virtually to discuss our goals, then broke up into teams to work on individual lessons before coming back together to review our progress. We hosted the curriculum in a public GitHub repository (https://github.com/umcarpentries/intro-curriculum-r) to facilitate collaborative work and peer review using issues, branches, and pull requests. Under this model, a team member created or edited content in a new branch to resolve an issue, then created a pull request and asked for review from another team member, who finally merged the changes into the default branch. GitHub pages automatically uses the default branch to build a website that allows us to host the polished curriculum (https://umcarpentries.org/intro-curriculum-r/). Our collaborative model ensured that at least two pairs of eyes viewed any changes before they could be included in the curriculum. This strategy helped us reduce mistakes and create better quality content.

**Figure 1:**
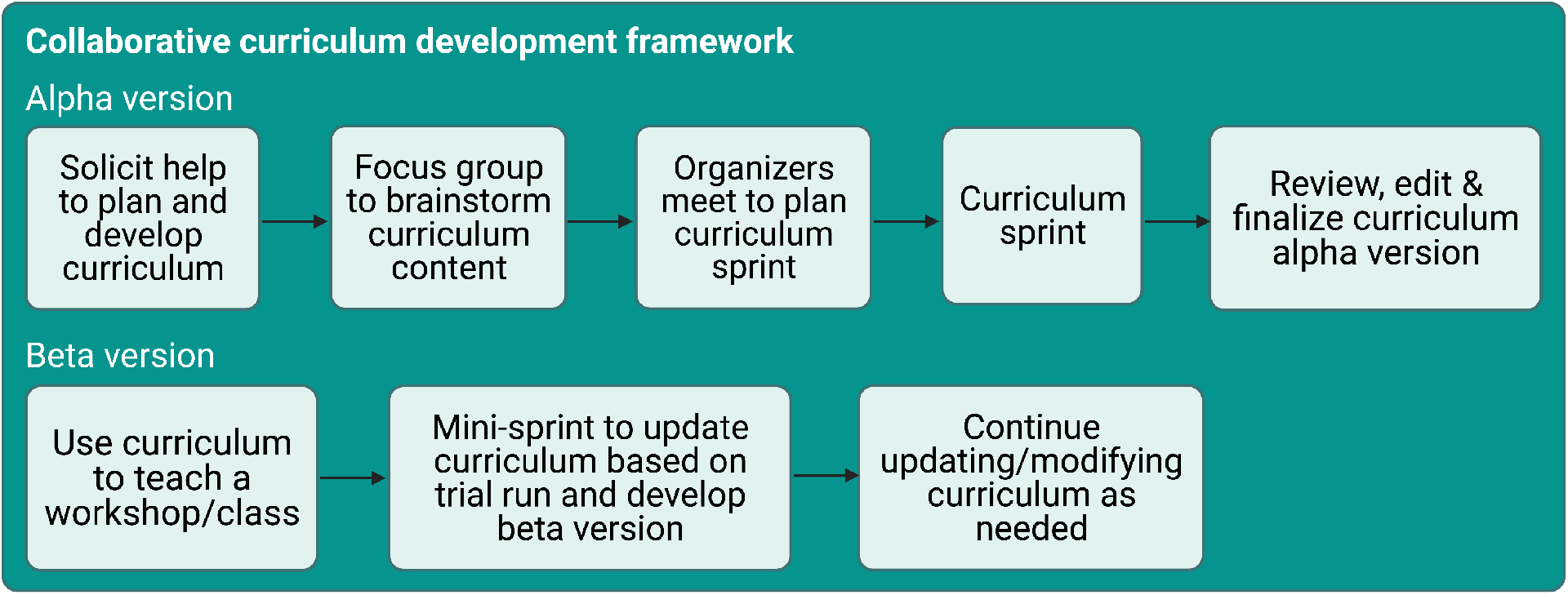
The curriculum development framework. Created with BioRender.com.

Following the sprint, contributors finalized edits and continued to review each others’ pull requests to complete the alpha version of our curriculum. Next, we hosted a workshop for instructors to pilot the curriculum. We collected feedback from the learners and instructors at the end of the pilot workshop and then held a smaller half-day sprint to revise the curriculum based on the feedback. Currently, our community members are continuously able to create issues, make edits, and review pull requests to keep refining the curriculum for future use. We are planning more workshops with new instructors who were not involved in the original curriculum development to gather their feedback.

## Curriculum

Our curriculum is tailored to people with no prior coding experience who want to learn how to use R programming for data analysis, visualization and the reporting of results (Figure 2). Not only do we aim to teach our learners the basics of performing empirical data analysis, we also seek to provide a rigorous framework for adhering to reproducible research principles that enable researchers to easily share their empirical work with others.

**Figure 2:**
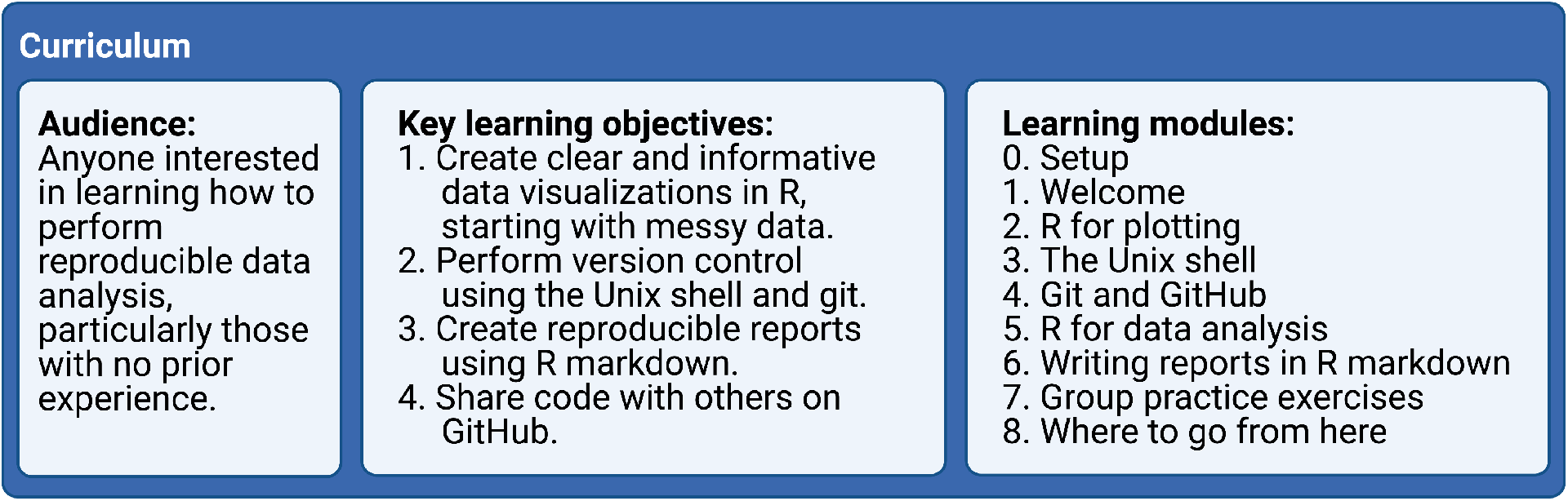
Curriculum overview. Created with BioRender.com.

### Learning Objectives

The key learning objectives for our curriculum are:

1. Create clear and informative data visualizations in R, starting with messy data.
2. Perform version control using the Unix shell and git.
3. Create reproducible reports using R Markdown.
4. Share code with others on GitHub.

We believe these skills provide learners with a solid foundation from which they can teach themselves any additional coding skills for future use.

### Course Content

Our curriculum consists of nine modules that cover software setup, data analysis and visualization in R, version control, sharing code, and writing reproducible reports (see below for more details). The R programming lessons take a “tidyverse first” approach (Robinson 2017) to effectively and efficiently teach learners powerful tools for plotting and data analysis. We also set an overall goal for the workshop to make the content substantively interesting and relatable to a wide audience regardless of their original academic discipline or professional practice. Specifically, we task our learners with producing a fictitious report to the United Nations that examines the relationship between gross domestic product (GDP), life expectancy, and CO_2_ emissions. The nine curriculum modules are:

0. Setup
1. Welcome
2. R for plotting (uses the tidyverse R packages (Wickham et al. 2019))
3. The Unix shell
4. Git and GitHub
5. R for data analysis (uses the tidyverse R packages (Wickham et al. 2019))
6. Writing reports in R Markdown (uses the rmarkdown R package (Xie, Allaire, and Grolemund 2018))
7. Group practice exercises
8. Where to go from here

Each lesson builds on the previous ones. The Unix shell, git, and GitHub are introduced using the files generated in the R for plotting lesson. The lesson content for subsequent modules is then intermittently committed and pushed to GitHub. The ‘Writing reports in R Markdown’ lesson combines all of the skills learned previously to produce a report that one could share with the United Nations. Next, learners put everything they have learned into practice by forming small groups and working on practice problems that cover the entire course content (“Integrating it all together: Paired exercise”). The workshop completes with a short module recapping everything that the curriculum covered as well as offering suggestions on how learners can continue to get help and keep learning once the workshop ends.

### Instructional Design

Our modules and teaching suggestions are developed in the style of Software Carpentry:

1. Each module contains learning objectives at the beginning of each lesson and a summary of key points at the end.
2. The five core modules (2 to 6) are designed to be taught via live coding of the content to learners. This is a central feature of Carpentries lessons, and we believe it is a great way to learn how to program. It requires learners to follow along and encounter errors that they must debug along the way, fostering additional questions about the course content. It also leads to instructors making mistakes and then demonstrating how to deal with them in an ad hoc and iterative manner.
3. We incorporate formative assessments in the form of short practice exercises throughout each lesson such that learners can practice what they have learned, while instructors can gauge learner understanding of the material.
4. We use the “sticky note” system for formative assessment, where learners indicate their progress on exercises and request help by using different colored sticky notes (Becker 2016; The Carpentries 2018a). At virtual workshops, we use Zoom reaction icons as virtual sticky notes, with the red X reaction to ask for help and the green checkmark to indicate that an exercise was successfully completed.
5. We have several helpers attend each workshop to address learner questions and technical issues.

We also incorporated a few additional key components into the curriculum:

1. Each lesson built off of previous lessons, with the goal of creating a final report that can be shared with others.
2. We structured the curriculum such that it could be taught through an in-person or virtual workshop. Virtual workshops are sometimes necessary, as during the COVID-19 pandemic, but are also useful to allow people from a variety of geographic locations to instruct and attend.
3. We not only required learners to install all software before the workshop (as The Carpentries also requires), but also asked them to run an example script that tests whether everything is installed correctly. To attend the workshop, learners were required to send screenshots of the script output to the workshop lead in advance. We withheld the login details for the workshop until we received the screenshot. This ensured that any installation issues could be addressed before the workshop began.
4. An extensive small group practice module towards the end of the workshop allowed learners to more independently practice the skills they have learned.
5. The workshop concluded with a recap of what was covered and resources available for learners to continue learning and getting help as their skills develop.

### Pilot Workshop

We piloted our curriculum during a virtual two-day Software Carpentry workshop. In line with The Carpentries recommendations (The Carpentries 2018b), we had four instructors and six helpers at the workshop to assist with learner questions and technical issues. We had thirty-nine learners of various skill levels from several different countries, all of whom provided very positive reviews of the workshop. To assess the effectiveness of the workshop, learners were asked to complete a pre- and post-workshop survey administered by the Carpentries. By the end of the workshop, learners on average felt more confident writing programs, using programming to work with data, overcoming problems while programming, and searching for answers to technical questions online (n = 14 survey respondents; see Figure 3). All attendees who filled out the post-workshop survey (n = 19) would recommend the workshop to others.

**Figure 3:**
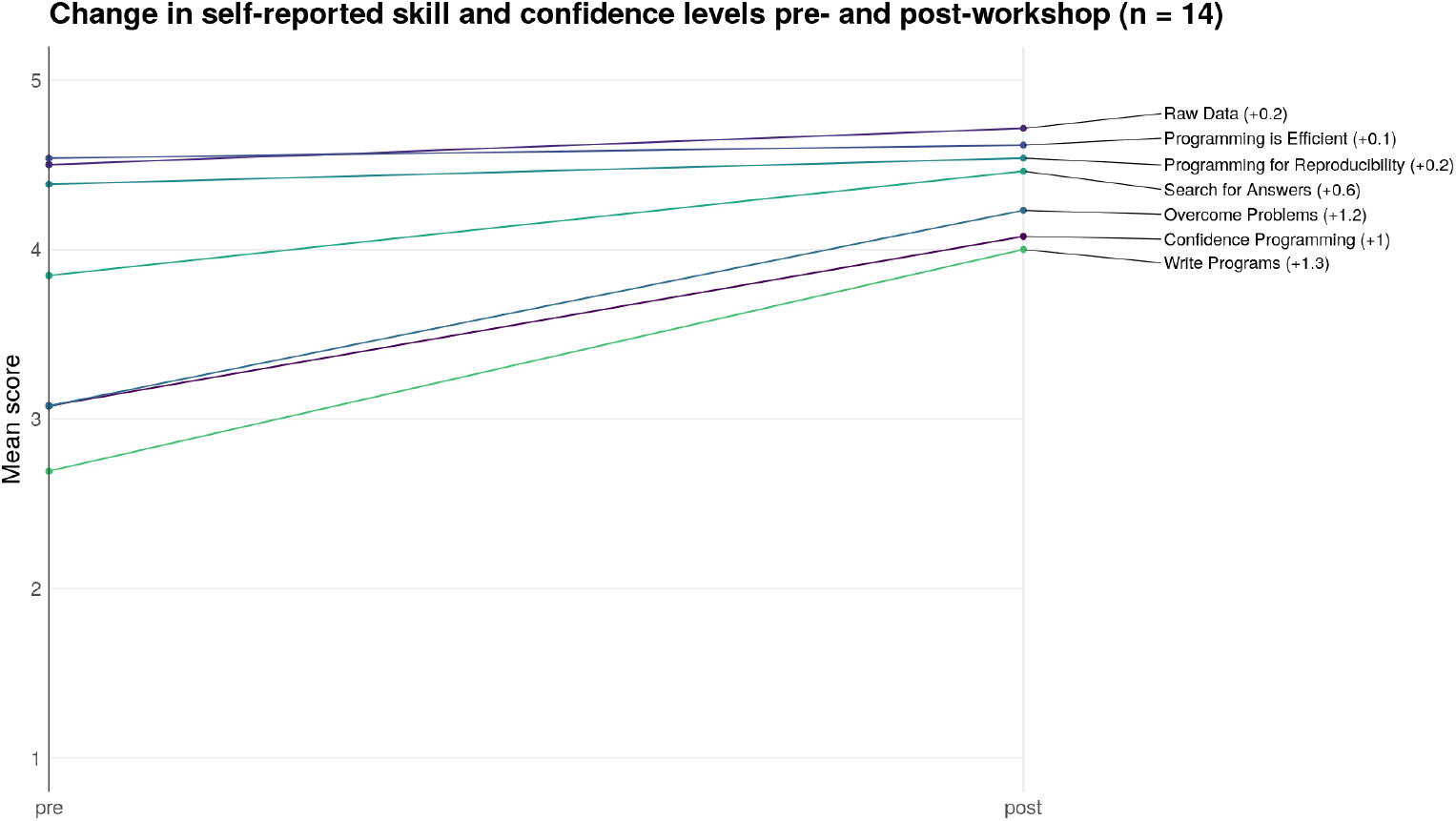
Pre- and post-workshop survey results.

### Virtual Workshop Reflection

We credit the success of our first virtual workshop in large part due to the curriculum structure and content, as well as the instructors and helpers involved. However, we also believe that the following helped make the workshop as smooth as possible:

1. We suggested that learners have Zoom and RStudio (or the Unix shell) open side-by-side on their computer to minimize toggling between different windows (Chen 2020).
2. We used Slack for communication among instructors and helpers, as well as between helpers and learners. Learners asked questions in a group Slack channel where helpers could respond. This allowed us to address the vast majority of learner questions and bugs quickly, clearly, and efficiently without disrupting the lesson or moving the learner to a Zoom breakout room. Furthermore, Slack worked much better than the Zoom chat as questions could be answered in threads, were preserved and visible to all learners regardless of whether they were connected to Zoom at the time, and didn’t get lost as easily.
3. Whenever a learner needed more help than was possible on Slack, a helper and the learner entered a Zoom breakout room together to troubleshoot. However, we tried to minimize this option as much as possible to prevent the learner from missing content covered in the main room.

## Acknowledgements

We thank The Carpentries organization for providing instructor training, workshop protocols, and the opensource Software Carpentry curriculum upon which this curriculum is based. We also thank them for allowing us to use the pre- and post-workshop survey results in this manuscript. The Carpentries is a fiscally sponsored project of Community Initiatives, a registered 501(c)3 non-profit organisation based in California, USA.

We are grateful to Victoria Alden and Scott Martin for assisting us in organizing and advertising our pilot workshop. We thank Shelly Johnson for volunteering as a helper at the workshop and contributing to the setup instructions. We also thank Bennet Fauber for contributing to the setup instructions.

We thank the learners who participated in the workshop, provided feedback, and completed the surveys.

## Funding

Salary support for PDS came from NIH grants R01CA215574 and U01AI124255. KLS received support from the NIH Training Program in Bioinformatics (T32 GM070449). ZL received support from the National Science Foundation Graduate Research Fellowship Program under Grant No. DGE 1256260. Any opinions, findings, and conclusions or recommendations expressed in this material are those of the authors and do not necessarily reflect the views of the National Science Foundation.

## Author Contributions

ZL and KLS contributed equally. ZL is first among the co-first authors because KLS threatened to reject all pull requests where ZL put KLS first. :)

PDS supervised the project. ZL and KLS organized the initial sprint, led the development of the curriculum, and drafted the manuscript. ZL, KLS, JK, and MML instructed at the first pilot workshop while CRA, JMA, ST, SKL, and CB assisted learners. All authors contributed to the development of the curriculum.

## Conflicts of Interest

None.

## Notes

https://github.com/umcarpentries/intro-curriculum-r

### Competing Interest Statement

The authors have declared no competing interest.

